# PEDF peptides rescue defects in neurite morphogenesis and intracellular calcium response in cortical neurons from mice exposed to valproic acid

**DOI:** 10.1101/2025.09.20.677502

**Authors:** Xiaonan Liu, Kazuhito Toyo-oka

## Abstract

Pigment epithelial-derived factor (PEDF) is a multifunctional protein produced predominantly by the retinal pigment epithelium and expressed in many tissues, including the brain, highlighting its participation in crucial processes, such as neuroprotection and angiogenesis. Some neurodevelopmental disorders, such as ASD, are characterized by neurodevelopmental abnormalities, including altered neurite formation, spine formation, and neuronal activities. Many efforts have been made to resolve NDDs, but until now, some symptoms remain untargeted. PEDF is involved in many steps of neurodevelopment. The treatment of PEDF peptide might improve the outcome of NDD symptoms by altering neuronal morphologies. We used PEDF peptides that contain different functional domains to study the effect of administering PEDF peptides on neuronal morphology in a prenatal valproic acid (VPA)-exposed mouse model. We identified that treatment with PEDF peptides rectified the abnormalities in neurite formation and spine formation in VPA-exposed cortical neurons. *In vitro* calcium imaging showed abnormalities in spontaneous activity in VPA-exposed cortical neurons. Treatment of a short PEDF peptide normalized the intracellular calcium response to the control level. Accordingly, PEDF peptides have the prospect of serving as potential treatments for patients with neurodevelopmental disorders, such as ASD.

## Introduction

Autism spectrum disorder (ASD) is a neuronal developmental disorder affecting approximately 1 in 34 children in the US as of 2022, according to the Centers for Disease Control (CDC). ASD is characterized by impaired communication and social interaction, repetitive behaviors, restrictive interests, and irritability. It was estimated that 40 to 80 percent of ASD is caused by genetic factors, while *de novo* mutations and epigenetic modulation by environmental factors also contribute to the cause ^1–3^. Due to the heterogeneity of ASD, it remains a challenge for basic research as well as treatment development for ASD. FDA has approved drugs for treating irritability, but therapies for the other three core symptoms are still under development. Currently, there are ongoing clinical trials for the development of treatments for various conditions in autism patients, including small molecules, peptides, medical devices, and behavioral interventions.

Pigment epithelial-derived factor (PEDF) is a glycoprotein with a molecular weight of about 50 kDa and is expressed in both glial cells and neurons in the central nervous system. PEDF is encoded by *SERPINF1* located on human 17p13.3, which has been found commonly disrupted in patients with Miller-Dieker Syndrome ^4,5^. Functionally, PEDF belongs to the non-inhibitory serpin family. It is known for its neuroprotective and antiangiogenic functions. Domains of PEDF, Val78-Thr121 (44-mer) and Asp44-Asn77 (34-mer), exert different functions putatively via variations in binding affinity to different cell surface receptors and the expression levels of these receptors in different cell types. The 44-mer binds to the PEDF receptor and exhibits neuroprotective functions in motor neurons, anti-inflammatory properties in the retina under diabetic retinopathy, and anti-apoptotic and anti-necrotic effects in hypoxic cardiomyocytes ^6–8^. On the other hand, the 34-mer peptide interacts with the Laminin receptor, resulting in antiangiogenic activity and inhibitory effect on progenitor cell proliferation in the liver ^9^. More receptors have been identified and studied for their functions, including cell-surface F1F0-ATP synthase and two transmembrane proteins, Plexin Domain Containing 1 (PLXDC1) and Plexin Domain Containing 2 (PLXDC2) ^10,11^. The receptors demonstrated significant functions in transducing PEDF signals, including neuroprotective effects on radial glial cells ^11,12^. Studies have been conducted to evaluate the effect of PEDF peptides in various model systems, exploring their treatment for disease conditions such as cancers, diabetic retinopathy, choroidal neovascularization, and bleomycin-induced pulmonary fibrosis ^7,13–15^.

The prenatal valproic acid (VPA)-exposed rodent model is a widely used model for studying ASD. Many studies have found that mice or rats exposed to VPA at the embryonic stage display ASD-like behaviors, such as reduced sociability, reduced exploratory behaviors, increased repetitive behaviors, and anxiety (reviewed by ^16^). Dendritic and axonal morphologies, as well as spine densities, are also changed in the mouse cortical neurons exposed to VPA before birth ^17–20^. A direct inhibition of the histone deacetylases (HDAC) and acetaldehyde dehydrogenase (ALDH1A1) by VPA at critical time points of development is known to be one of the main mechanisms contributing to the autistic phenotypes in VPA prenatal-exposure models ^21–24^. Indeed, histone acetylation regulates the expression of the genes associated with ASD, such as *PTEN* and *RELN* ^25,26^. Notably, *SERPINF1*, the gene that codes for PEDF, is also under the control of HDAC ^27^. Our lab identified a novel role of PEDF in mouse cortical neuronal development ^28^. We found that PEDF regulated neurite formation during cortical development. Thus, PEDF peptides could be beneficial in treating disorders like ASD, by which abnormalities in neuromorphogenesis would be ameliorated.

In the present study, we used cultured cortical neurons from VPA-exposed embryos to investigate the effect of PEDF peptides on neurite formation, spine formation, and spontaneous calcium response. We showed that the 44-mer PEDF was able to correct the reduced neurite length in the longest neurite in the VPA-exposed neuron at DIV2. Moreover, the 44-mer PEDF rectified the reduced spine density and enlarged mushroom spine head in the VPA-exposed neurons at DIV16. Analysis of spontaneous calcium response showed that the 44-mer PEDF also rectified the abnormal calcium response. The findings may advance our understanding of PEDF in neuronal morphogenesis and facilitate further studies of the peptide as a potential treatment for ASD.

## Results

### PEDF peptides promoted neurite formation in neurons from VPA-exposed mice

To test the effect of PEDF peptides on neurite formation, we cultured cortical neurons prepared from control or prenatally VPA-exposed mice with synthetic PEDF peptides for two days (Fig. 1). Then, we analyzed the length of the longest neurite, the length of shorter neurites, and the number of neurites.

**Figure 1.**
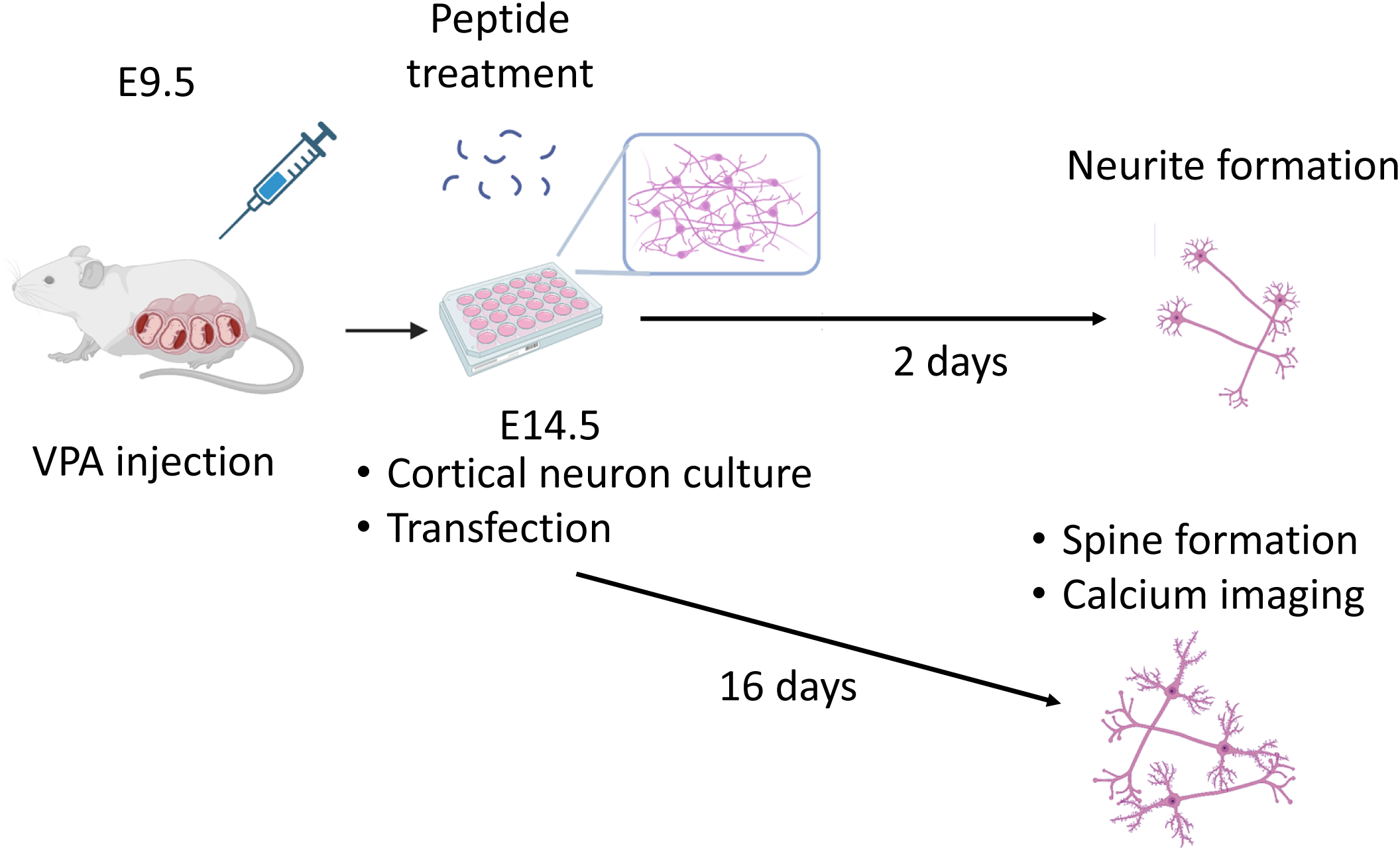
Illustration of the experiment design for studying *in vitro* neurite formation using maternal VPA-exposed mice. Embryos were exposed to VPA at E9.5. Cortical neurons from these embryos were then dissected and cultured with vehicles or peptides supplied in the culture media at E14.5. DIV2 neurons were used to study neurite formation, while DIV16 neurons were obtained to study dendritic spine formation and spontaneous calcium response.

We found that the longest neurite in the neurons from VPA-exposed mice, which is likely to become the axon in each cell, was significantly shorter than that of the controls. After treating the VPA-exposed cells with the peptides, the neurite length was corrected by the 44-mer and the 18-mer, but not by the 34-mer (Fig. 2A and B). The length of shorter neurites, which are likely to become dendrites, was not significantly affected by the prenatal VPA exposure, as well as the treatment by the peptides (Fig. 2C). Neurons treated with VPA showed fewer neurites compared to the control group (Fig. 2D). The treatment with the 44-mer, 34-mer, and 18-mer peptides successfully rescued the defects in neurite number observed in neurons from VPA-exposed mice (Fig. 2D).

**Figure 2.**
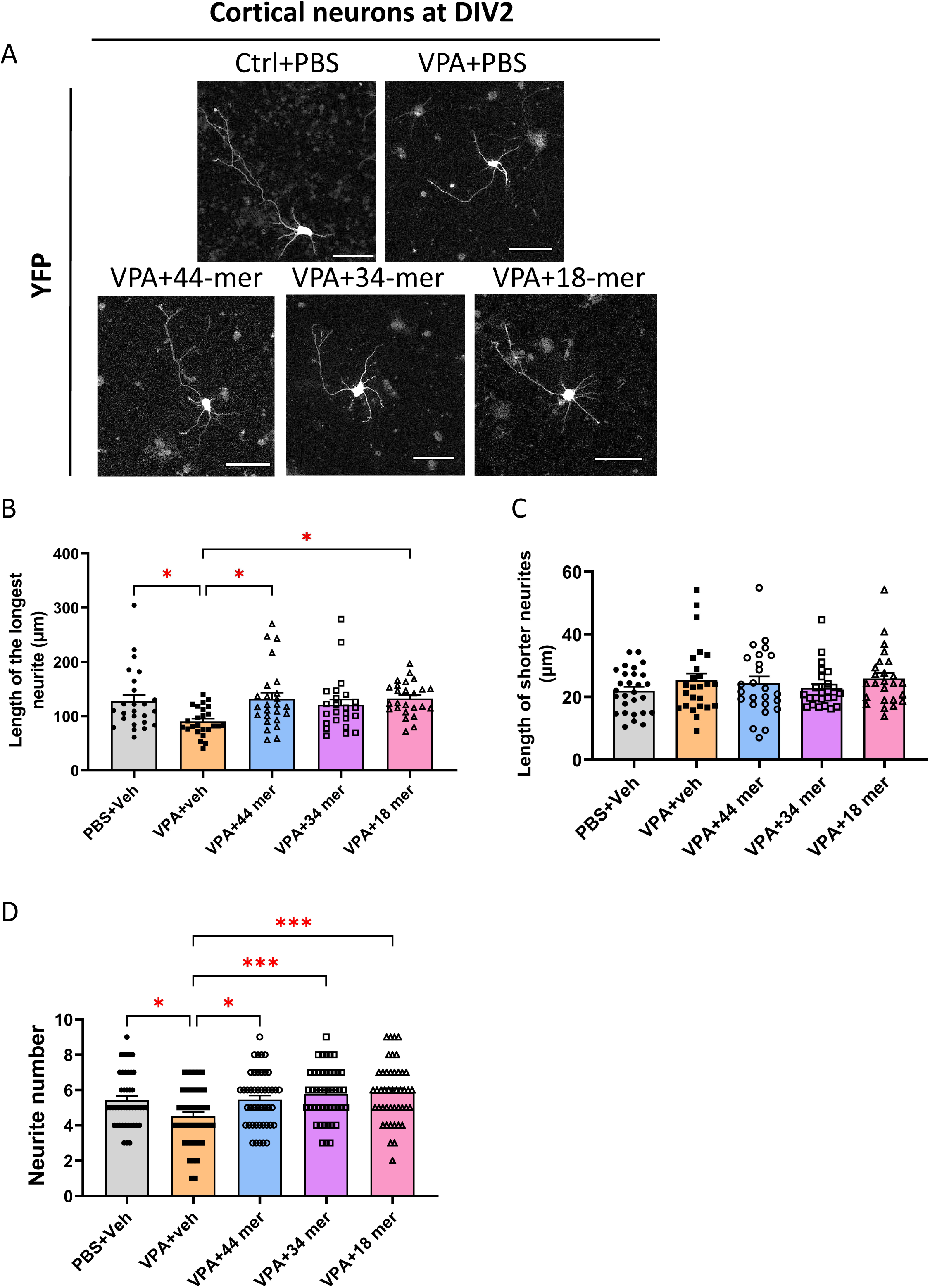
PEDF peptide treatments altered neurite formation in DIV2 neurons from mice with prenatal VPA exposure. A) Representative images of DIV2 neurons expressing YFP from different treatment groups, including control neurons treated with vehicle (PBS), neurons from prenatal VPA-exposed offspring treated with vehicle, neurons from VPA-exposed offspring treated with 44-mer PEDF peptides, neurons from VPA-exposed offspring treated with 34-mer peptides, and neurons from VPA-exposed offspring treated with 18-mer peptides. Scale bar=50μm. B) Two-way ANOVA determined significant differences in the length of the longest neurites between the six groups (F(4,95)=3.566; ** P<0.01; n=24-25 cells per group). Tukey’s multiple comparison showed a reduction in the longest neurite length in the VPA model with PBS treatment (90.15 ± 5.214, n=25 cells) compared to the control with PBS treatment (127.5 ± 11.31, n=25, * P<0.05), and significant increase by 44-mer PEDF (131.7 ± 11.42, n=25 cells; * P<0.05), and 18-mer PEDF treatments (132.4 ± 6.222, n=25 cells; * P<0.05), but not by 34-mer PEDF (120.4 ± 10.22, n=24, P=0.1511 treatment. No difference was found in the length of the longest neurite in Ctrl neurons with PBS and VPA neurons with 44-mer (P=0.9977), or 18-mer treatment (P=0.9956). C) Two-way ANOVA determined no significant change in the average length of shorter neurites (F(4,96)=0.7482; P=0.5616; n=25-28 cells per group). Multiple comparisons showed no statistical significance in the six groups. D) Two-way ANOVA showed a significant difference in the neurite number (F(4,174)=5.734; *** P<0.001; n=45 cells per group). Tukey’s multiple comparison showed a significant difference between the Ctrl neurons with PBS treatment (5.444 ± 0.2282) and VPA neurons with PBS treatment (4.511 ± 0.2413; * P<0.05). A significant increase was found between the VPA-exposed neurons with PBS treatment and VPA-exposed neurons with 44-mer treatment (5.467 ± 0.2327; * P<0.05), 34-mer treatment (5.778 ± 0.2222; ** P<0.01), and 18-mer treatment (5.867 ± 0.2494; *** P<0.001).

### PEDF peptide altered the composition of dendritic spines in cortical neurons prepared from VPA-exposed mice

Dendritic spine density and structure are often altered in ASD patients and animal models, which is believed to contribute to autistic-like behaviors, such as repetitive behavior, impaired cognitive function, and social disability ^29–31^. A key question to answer for many potential ASD therapies is whether the treatment can correct the abnormal dendritic spine morphology in ASD. We analyzed the effect of the peptides on dendritic spine structures.

We cultured primary cortical neurons from control or VPA-exposed embryos for 16 days *in vitro* (DIV16), which corresponds to the time period for dendritic spine formation and maturation *in vitro* (Fig. 1) ^32^. PBS or a 44-mer peptide was added to the culture media at the beginning of the culturing process. The cells were transfected with the plasmid coding for YFP at DIV6 to identify and analyze neuromorphology.

We found that VPA prenatal exposure at E9.5 significantly reduced the dendritic spine densities in cortical neurons (Fig. 3A and B). Treatment with the 44-mer PEDF, however, rescued the density to the control level (Fig. 3A and B).

**Figure 3.**
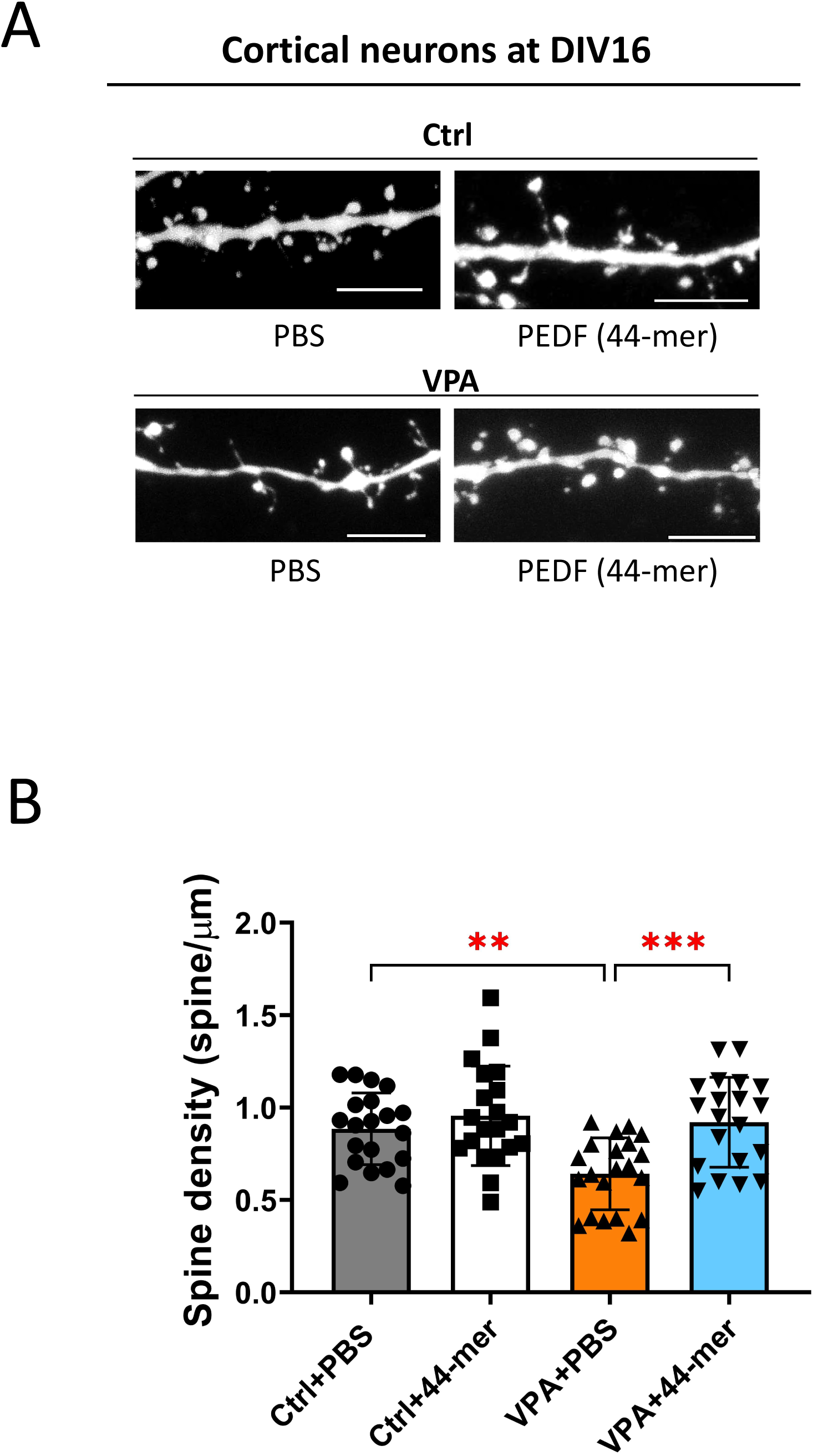
PEDF peptide treatment altered dendritic spine density in DIV16 neurons from mice with prenatal VPA exposure. A) Representative figures of dendritic spine morphologies in neurons at DIV16. Scale bar=5μm. B) Quantification of dendritic spine density in control cells treated with PBS or 44-mer, and VPA-exposed cells treated with PBS or 44-mer peptides. Two-way ANOVA showed a significant difference between the four groups (F(3,57)=10.46; P<0.0001; n=20-21 cells per group). Tukey’s multiple comparisons determined a statistically significant reduction in spine density in neurons from the VPA model with PBS treatment (0.6421 ± 0.04246) compared to the Ctrl with PBS treatment (0.8853 ± 0.04342; ** P<0.01). A statistically significant increase was identified in the neurons from the VPA model with 44-mer treatment (0.9209 ± 0.05433) compared to neurons from the VPA model with PBS treatment (0.6421 ± 0.04246) (*** P<0.001).

Dendritic spines were canonically classified based on their shapes into filopodia, long thin, thin, stubby, mushroom, and branched, which reflect the signal transduction properties (Fig. 4A). Mushroom spines and branched spines are usually known as mature spines that hold the strongest synaptic properties. Thin spines are new spines that have the potential to grow into mature spines, which are also called learning spines. Stubby spines are immature spines that hold less synaptic material than mature spines. We found that VPA-exposed neurons showed a significantly decreased number of thin spines compared to control neurons, while treatment with the 44-mer peptides ameliorated this abnormality (Fig. 4B). The densities of other spine types were not significantly altered. Although the mushroom spine density was not changed in the VPA-exposed neurons, when we analyzed the mushroom spine head width, which reflects the synaptic strength of the mushroom spines, VPA-exposed neurons had larger mushroom spines, while the treatment of the 44-mer rescued this abnormality (Fig. 4C). The results suggested that in the VPA-exposed neurons the treatment with 44-mer peptides rescued the spine density, spine maturation and synaptic function.

**Figure 4.**
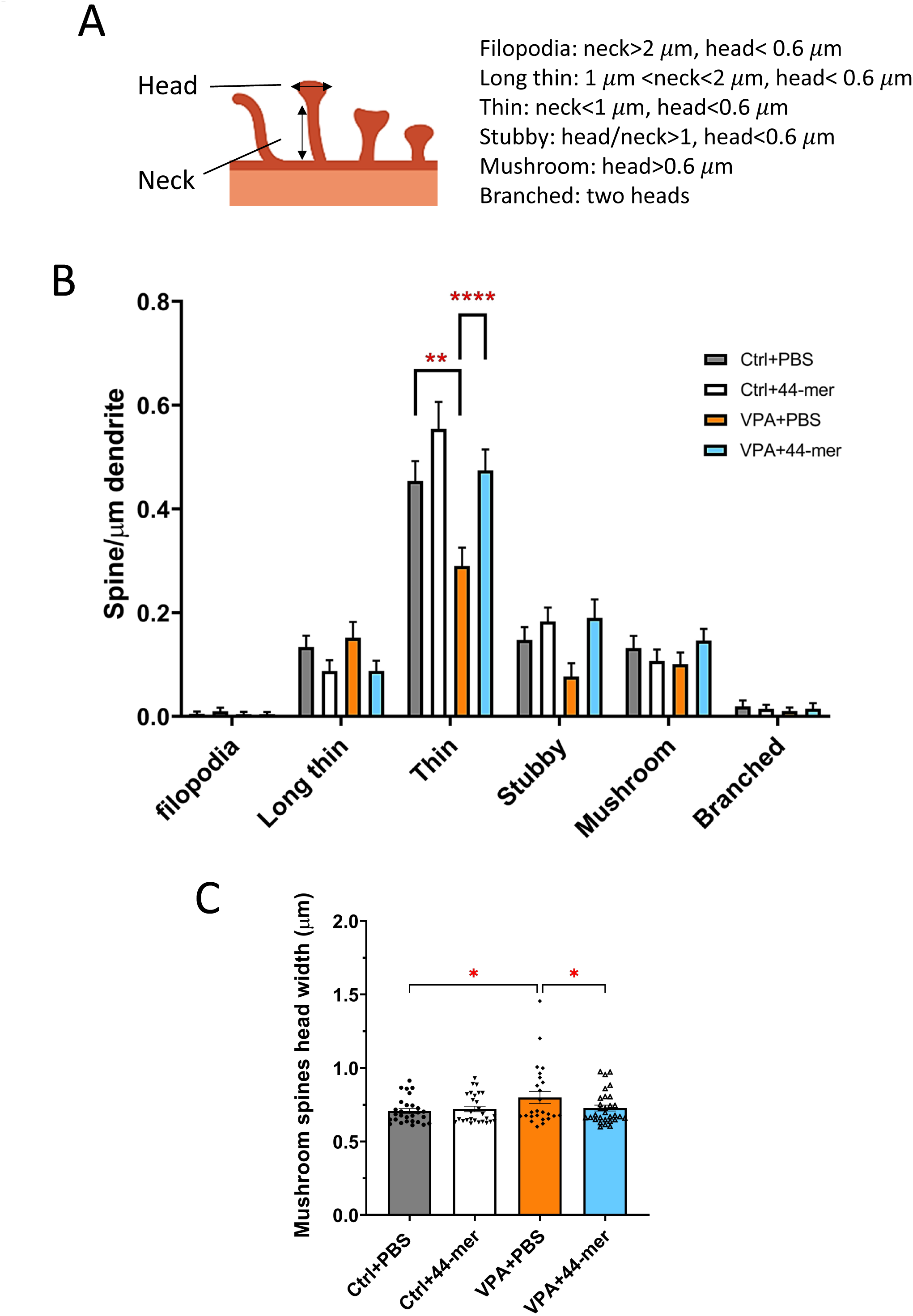
PEDF peptide treatment altered dendritic spine morphologies in DIV16 neurons from mice with prenatal VPA exposure. A) Illustration of the quantitative criteria of spine morphology classes. Adapted from Risher et al. ^32^. B) Quantification of the density per micron of the six spine types, respectively. A three-way ANOVA revealed a significant difference among the treatments in the different mice across the spine types (F(1,76)=4.434; *P<0.05). Tukey’s multiple comparison showed reduced density in the thin spines in the neurons from VPA-exposed mice with PBS treatment (0.29 ± 0.035; n=20 cells) compared to the neurons from the control mice with PBS treatment (0.454 ± 0.039; N=20 cells) (** P<0.01). 44-mer treatment significantly increased the thin spine density in the VPA model (0.474 ±0.04, n=20); **** P<0.0001), and to the same level as neurons from Ctrl mice treated with PBS (P>0.9999). No significant difference was found in other spine types between neurons from control mice and those from VPA mice. C) Quantification of the head width of mushroom spines. A two-way ANOVA determined a significant difference in the width between groups (F(3,77)=3.910; n=25-30 cells per group: * P<0.05). Tukey’s multiple comparison showed the head width is larger in the neurons from the VPA model with PBS treatment (0.7993 ± 0.04112; n=25 spines) compared to the neurons from the Ctrl with PBS treatment (0.7075 ± 0.01649; n=28 spines) (* P<0.05). The width of the mushroom spine is significantly reduced by applying 44-mer peptides to the neurons from the VPA model (0.7272 ± 0.01987; n=30 spines) (* P<0.05). No difference was found between the neurons from Ctrl mice treated with PBS and the neurons from the VPA model mice treated with 44-mer (P=0.9864).

### The treatment with the 44-mer PEDF peptide rescued abnormal spontaneous calcium response in neurons from VPA-exposed mice

We found structural abnormalities in the dendritic spines of DIV16 cortical neurons from VPA-exposed mice compared to those from control mice (Figs 3 and 4). The altered dendritic spine structure may result in a change in neuronal activity. To study this at DIV16, we performed spontaneous calcium response recordings in cortical neurons from VPA-exposed or control mice with or without treatment with the 44-mer PEDF peptide (Fig. 5).

**Figure 5.**
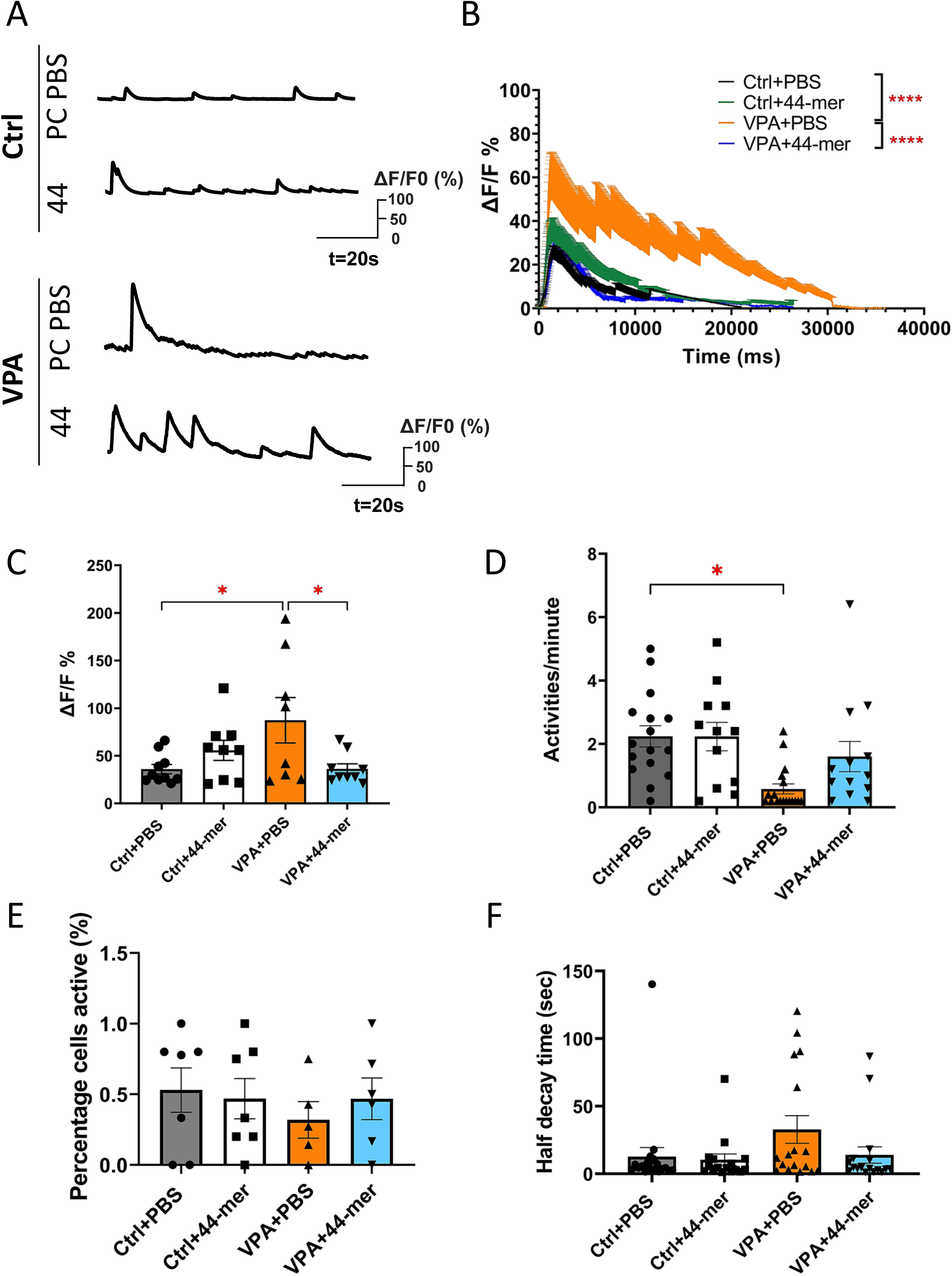
PEDF peptide treatment altered spontaneous calcium response in DIV16 neurons from mice with prenatal VPA exposure. PEDF peptide treatment altered spontaneous calcium response in DIV16 neurons from mice with prenatal VPA exposure. A) Representative tracings of the recording of fluorescence change over time. B) Summarized tracing of spontaneous calcium activities with the above error bars. Black (Ctrl neurons treated with PBS), green (Ctrl neurons treated with 44-mer), orange (VPA-exposed neurons treated with PBS), blue (VPA-exposed neurons treated with 44-mer). Two-way ANOVA showed a significant difference in the tracings between the groups (F(3,13060)=556.9, **** P<0.0001; n=9-11 activities per group). Tukey’s multiple comparisons showed there was a significant difference in the curves of the Ctrl neurons treated with PBS and VPA-exposed neurons treated with PBS (**** P<0.0001), difference in the curves of the VPA-exposed neurons treated with PBS and the VPA-exposed neurons treated with 44-mer (**** P<0.0001), while no difference between the curves of Control cells treated with PBS and VPA-exposed cells treated with 44-mer (P=0.5376). C) Quantification of fluorescence change ΔF/F0% of spontaneous calcium activities over the 23 ± 3.2% threshold determined by Chen et al. ^55^. A two-way ANOVA showed a significant difference in the ΔF/F0% between groups (F(3,22)=4.158, * P<0.05). Tukey’s multiple comparisons showed a significant increase in the fluorescence changes in the neurons from the VPA model treated with PBS (87.26 ± 23.84; n=8 activities) and the neurons from Ctrl model treated with PBS (35.97 ± 4.957; n=10 activities) (* P<0.05). Applying 44-mer to the neurons from the VPA model (36.22 ± 5.319, n=9 activities) significantly reduced ΔF/F0% compared to the neuron from the VPA model with PBS treatment (P=0.0277), while ΔF/F0% in the 44-mer treated VPA-exposed neurons did not changed compared to the ΔF/F0% in the Ctrl cells treated with PBS (P=0.9997). D) Quantification of spontaneous calcium response per minute. A two-way ANOVA determined there was a significant difference between the groups (F(3,38)=4.435; ** P<0.01). Tukey’s multiple comparisons identified a substantial reduction in the activities per minute in the neurons from the VPA model treated with PBS (0.5778 ± 0.1575, n=18 cells), compared to the neurons from the Ctrl model treated with PBS (2.238 ± 0.3373, n=16 cells) (* P<0.05). The 44-mer treatment in the neurons from the VPA model (1.6 ± 0.4788, n=13 cells) showed a trend of rescue compared to the PBS-treated neurons from the VPA model (* P<0.05). E) Quantification of the percentage of cells that had activity during the recording of each experiment. A Two-way ANOVA showed no difference between the groups (F(3,15=0.2188, P=0.8818). F) Quantification of half decay time in each activity. A two-way ANOVA showed no difference between the groups (F(3,47)=2.259, P=0.0938).

First, the spontaneous calcium response in the DIV16 neuron from VPA-exposed mice showed a different tracing pattern compared to that of neurons from control mice (Fig. 5A and B). VPA-exposed neurons treated with 44-mer PEDF had a similar curve as the control mice (Fig. 5B). Comparing the fluorescence changes, we found neurons from VPA-exposed mice had higher amplitude than the control (Fig. 5C). Applying the 44-mer PEDF to neurons exposed to VPA decreased the fluorescence change to the same level as in the control (Fig. 5C). The frequency of spontaneous calcium response was lower in the neurons from VPA-exposed mice compared to the control (Fig. 5D). While treatment with 44 PEDF in the VPA-exposed group showed a trend of rescue, it was not statistically significant (Fig. 5D). The percentage of cells that are active during the period of recording and the half time of decay are not changed in all the groups (Figs. 5E and F). Overall, the results showed that the 44-mer PEDF peptide rectified the increased spontaneous calcium amplitude and partially rescued the reduced frequency in the neurons from prenatal VPA-exposed mice.

## Discussion

Our study showed that treatment with PEDF peptides rescued neurite morphologies, spine structures, and neuronal activity in an ASD model *in vitro*. Neurons from VPA-exposed mice had a reduced length of the longest neurite at DIV2. The length of the longest neurite in the VPA-exposed neurons was rescued by 44-mer and 18-mer PEDF peptides. On the other hand, the length of shorter neurites did not change with VPA exposure. The number of neurites from the soma was reduced in VPA-exposed neurons, and treatment with the peptides, 44-mer. 34-mer, and 18-mer, rescued these defects. We analyzed *in vitro* dendritic spine morphology at DIV16 following treatment with a 44-mer PEDF peptide. The reduced spine density in the neurons from VPA-exposed mice was mitigated by the 44-mer. Mushroom spines are known as mature spines and have a predominant function in synaptic connections. In neurons from VPA-exposed mice, mushroom spine head widths were increased compared to those in control neurons, while treatment with 44-mer ameliorated this increase in width. Moreover, to further understand the effect of 44-mer on neurons from VPA-exposed mice, we performed in vitro calcium imaging to investigate spontaneous calcium activity. Calcium imaging revealed an increase in fluorescence change following prenatal VPA exposure. The administration of the 44-mer corrected the fluorescence change to the control level. An impairment of the frequency was found in VPA-exposed neurons, and this was also ameliorated by 44-mer treatment. Thus, these results showed that the treatment with the 44-mer peptide was able to correct the abnormal spontaneous calcium response in DIV16 neurons from prenatal VPA-exposed mice. Collectively, the experiments indicated that the 44-mer PEDF peptide treatment in the ASD mouse model rescued abnormalities in neurite morphology at the early neuronal morphogenesis stage and in dendritic spine structure at the later stage, as well as spontaneous calcium response.

Prenatal exposure to VPA is a widely studied model for ASD. VPA exposure affects neurogenesis, neurite formation, and spine morphology in the offspring, and alters neuronal sensory functions in multiple brain regions, contributing to behavioral changes ^19,33–36^. The prenatal VPA exposure model is widely used for studying ASD; however, the resulting abnormalities or phenotypes from *in utero* VPA exposure differ by the gestation date of exposure, the amount of VPA, the timing of observation, the type of neuron or brain region analyzed, as well as the genetic background of the mouse used for the experiment. It is generally accepted that VPA-exposed cortical neurons display delayed neurite outgrowth, with a decrease in the length of both the longest and shortest neurites. However, conclusions regarding spine morphology vary across studies ^16^. In our study, we analyzed neuronal morphogenesis in VPA-exposed neurons and found results similar to those reported in previous studies.

After prenatal VPA exposure, rodents exhibit altered neuronal activity in multiple brain regions and across different types of neurons ^37–39^. In our *in vitro* calcium imaging, we found abnormal spontaneous calcium activity in the DIV16 mouse cortical neurons from maternally VPA-exposed offspring as well. An increased fluorescence change of the calcium indicator and a reduced frequency were observed in the neurons from the VPA model during recording. Treatment with the 44-mer PEDF corrected both activity changes from the levels found in the VPA-exposed neurons to the control levels. The altered activities may be associated with the morphological changes observed in DIV2 and DIV16. The spontaneous calcium response we recorded in the cultured cortical neurons may arise from a mixed population of excitatory and inhibitory neurons. However, approximately 60% to 80% of neurons in the cerebral cortex at this time point are excitatory ^40^. Future studies could investigate the neuronal activities in specific neuronal populations to unveil more precise effects of the PEDF peptide on neuronal activity.

ASD affects approximately 60 percent more males than females ^41^. VPA *in utero* exposure resulted in autistic phenotypes also displaying a sex difference ^42^. In behavioral studies, reduced sociability, impaired cognition, and dysfunctional memory were observed only in the male offspring ^43–45^. At a molecular level, maternal VPA exposure differentially modulates DNA and RNA expression in males and females ^17,46^. In the current experiments, we used neurons from a combination of male and female offspring. Studying the effects of PEDF peptides in males and females separately might give more insight into the peptide effects on neurodevelopmental abnormalities caused by prenatal VPA exposure.

Previously, our lab identified that PEDF regulates neuromorphogenesis in cortical neurons. PEDF knockdown in cortical neurons causes reduced primary neurite number, shortened dendritic length, and decreased dendritic spine density ^28^. This previous study also showed that one of the PEDF receptors, non-integrin 37/67-kDa laminin receptor/ribosomal protein SA (RPSA), is responsible for transmitting neuromorphogenesis signaling. Currently, six PEDF receptors have been found, including RPSA, PEDF receptor (PEDF-R), catalytic β subunit of F1F0-ATP synthase, low-density lipoprotein receptor-related protein-6, PLXDC1, and PLXDC2 ^11,12,47–50^. Studies by other groups proposed that short peptide fragments derived from PEDF may act on different receptors on the cellular membrane (^51^). Thereby, differential downstream pathways may be evoked by the peptides. In the present study, we found that the three peptides exhibited distinct functions in neurite formation in the VPA model. The 44-mer and 18-mer rescued the length of the longest neurite and the number of neurites. The 34-mer rescued the number of neurites. Although we focused on the 44-mer in spine formation and neural activities, the effects of 34-mer and 18-mer should be further investigated. Furthermore, the molecules and signaling pathways regulated by administering the peptides could be investigated by using multi-omics analysis, which might reveal molecular mechanisms behind the PEDF peptide treatment.

The mechanism of prenatal VPA exposure-induced ASD is reported to be that VPA acts as an inhibitor for histone deacetylases (HDACs). HDACs modify the expression of many genes, including multiple ASD-risk genes such as *NLGN1*, *SHANK2*, *SHANK3*, and *CNTNAP2* ^26^. PEDF is found to be affected by HDAC inhibition ^52^, suggesting that PEDF expression may be under the regulation of HDACs. Therefore, there is a possibility that PEDF peptide treatment may be limited to the VPA exposure model and other models for ASD related to HDAC functions. Therefore, the effects of PEDF peptides on other types of ASD models should be further investigated.

The present study suggested a potential path to develop PEDF peptides for treating neurodevelopmental disorders. We have shown that neurite outgrowth, spine morphology, and spontaneous calcium response were affected in the VPA neurons and corrected by the PEDF peptide. In the future, a more thorough understanding of PEDF peptides will enhance our understanding of their effects.

## Material and Methods

### Plasmids and chemicals

Mouse PEDF peptides were synthesized by Genemed Synthesis, Inc. The sequences of PEDF peptides are as follows: 44-mer (78-121), VLLSPLSVATALSALSLGAEHRTESVIHRALYYDLITNPDIHST, 34- mer (44-77), DPFFKVPVNKLAAAVSNFGYDLYRLRSSASPTGN, 18-mer (60-77), NFGYDLYRLRSSASPTGN. pCAG-YFP plasmid was gifted from Connie Cepko (Addgene plasmid # 11180; http://n2t.net/addgene:11180) ^53^.

### Prenatal VPA exposure

The C57BL/6 mouse was used for the VPA injection. Female mice were mated, and embryonic days are timed by checking the vaginal copulatory plugs the next morning. VPA (500mg/kg) was intraperitoneally injected at E9.5 to induce an autism model as previously described ^19,54^.

### Primary cortical neuron culture and peptide treatment

Pregnant dams at E14.5 were sacrificed, and embryonic cortices were dissected for primary neurons as previously done. Briefly, the cortices were treated with 0.01% trypsin to achieve a single-cell suspension. 50 mg/ml BSA was used to inactivate the trypsin. For short-term culture, cells were plated on coverslips and fixed at two days after plating. For long-term culture, cells were plated on coverslips, and culture media were replaced every three days with fresh culture media supplemented with AraC.

The synthetic PEDF peptides were added to the culture media after cell seeding and remained present throughout the cell culture period. For each of the PEDF peptides, a final concentration of 5μM was used for the experiments ^28^. Ca2+/Mg2+-free Dulbecco’s PBS (D-PBS, Genesee Sci) was used as a vehicle control since the peptides were dissolved in D-PBS.

### Transfection

For nucleofection, cells were transfected as the manufacturer instructed (Mirus Bio). 100 μl of Mirus reagent and 10 μg of plasmids were used for nucleofection. The mixture was transferred to the cuvette (Ingenio). Nucleofection was performed using the Amaxa Nucleofector II system. After electroporation, the cell mixture was quickly transferred into a tube containing 500μl of culture media.

For chemical transfection with Lipofectamine LTX and PLUS reagent, cells were transfected at DIV6 according to the manufacturer’s instructions (Thermo Fisher Scientific). Briefly, 0.5μg of DNA was added to 100μl Neurobasal media, then 0.5μl of PLUS reagent and 1.25μl of LTX reagent were added; the mixture was incubated at room temperature for 30 min before being applied to the cell culture.

### *In vitro* calcium imaging and analysis

In vitro calcium imaging experiments were conducted at DIV16. GCaMP6s was used as a calcium indicator and was transfected into the cells at DIV6. An inverted fluorescence microscope (Zeiss) with a 20X objective and a temperature-controlled chamber maintained at 37°C was used for time-lapse imaging. The cell culture medium was replaced with DMEM/F12 without phenol red for the recording. Images were captured every 35 msec for about 5 minutes. The fluorescence intensity of the soma area was analyzed using Zen 2 Pro analysis software (Zeiss, 2011). The percentage fluorescence change of each time point was calculated as Δ*F*/*F* × 100, where F is the baseline fluorescence intensity, Δ*F* is the fluorescence intensity of each time point minus *F*, as previously described ^55^. A fluorescence change in response to one action potential was determined by the fluorescence change over 23±3.2% ^55^.

### The Analyses of Neuronal Morphology

Images were captured using a confocal microscope (Leica SP8), and all analyses were performed in Fiji ImageJ. For neurite formation analyses, neurites were traced manually to measure length and number. To analyze dendritic spine morphology, we adopted the classification method described by Risher et al. ^32^. Images were taken using a 63x objective lens on a confocal microscope (Leica SP8) with z-stacks to record the structures on neuronal secondary and tertiary dendrites. The length of the dendritic spine neck and the width of the dendritic spine head were measured using z-projection images. Per dendritic spine with >=0.6 μm-head was considered as mushroom spine, with < 0.6 μm-head and >2 μm-neck was identified as filopodia, with <0.6 μm-head and 1-2 μm-neck was identified as long thin spine, with <1 μm-neck and a head width less than neck was found to be thin spine, and with <0.6 μm-head and a ratio of head to neck >1 was identified as stubby spine ^32^.

For dendritic spine density, the number of dendritic spines about 10 μm was counted in each measurement. Dendritic spine per μm was calculated by dividing the number of spines by the length of the fragment of dendrite in each measurement.

### Statistics

Quantitative data were analyzed using Prism (GraphPad). The data were analyzed using two-way ANOVA, and three-way ANOVA, as appropriate. Values were represented as the mean (SEM). Results were determined to be statistically significant if the P value was <0.05, * P<0.05, ** P<0.01, *** P<0.001, and **** P<0.0001.

## Acknowledgments

This work has been supported by R01 grant from NINDS (NS096098), R21 grant from NIMH (R21MH132839) and autism grant from the Department of Defense (AR220067) (to K.T.).

## Author Contributions

X. L. performed all experiments and analyses. She also performed all mouse breeding and maintenance and wrote the manuscript. K.T. supervised all experiments and analyses and finalized the manuscript.

## References

1 Chaste, P. & Leboyer, M. Autism risk factors: genes, environment, and gene-environment interactions. Dialogues Clin Neurosci 14, 281–292 (2012).

2 Wong, C. C. et al. Methylomic analysis of monozygotic twins discordant for autism spectrum disorder and related behavioural traits. Mol Psychiatry 19, 495–503 (2014).

3 Duffney, L. J. et al. Epigenetics and autism spectrum disorder: A report of an autism case with mutation in H1 linker histone HIST1H1E and literature review. Am J Med Genet B Neuropsychiatr Genet 177, 426–433 (2018).

4 Homs, A. et al. Genetic and epigenetic methylation defects and implication of the ERMN gene in autism spectrum disorders. Transl Psychiatry 6, e855–e855 (2016).

5 Blazejewski, S. M., Bennison, S. A., Smith, T. H. & Toyo-Oka, K. Neurodevelopmental Genetic Diseases Associated With Microdeletions and Microduplications of Chromosome 17p13.3. Front Genet 9, 80–80 (2018).

6 Bilak, M. M. et al. Identification of the Neuroprotective Molecular Region of Pigment Epithelium-Derived Factor and Its Binding Sites on Motor Neurons. J Neurosci 22, 9378 (2002).

7 Liu, Y. et al. Pigment epithelium-derived factor (PEDF) peptide eye drops reduce inflammation, cell death and vascular leakage in diabetic retinopathy in Ins2(Akita) mice. Mol Med 18, 1387–1401 (2012).

8 Gao, X. et al. PEDF and PEDF-derived peptide 44mer protect cardiomyocytes against hypoxia-induced apoptosis and necroptosis via anti-oxidative effect. Sci Rep 4, 5637 (2014).

9 Protiva, P. et al. Pigment Epithelium-Derived Factor (PEDF) Inhibits Wnt/β-catenin Signaling in the Liver. Cell Mol Gastroenterol Hepatol 1, 535–549.e514 (2015).

10 Deshpande, M., Notari, L., Subramanian, P., Notario, V. & Becerra, S. P. Inhibition of tumor cell surface ATP synthesis by pigment epithelium-derived factor: implications for antitumor activity. Int J Oncol 41, 219–227 (2012).

11 Cheng, G. et al. Identification of PLXDC1 and PLXDC2 as the transmembrane receptors for the multifunctional factor PEDF. eLife 3, e05401 (2014).

12 Notari, L. et al. Pigment epithelium-derived factor binds to cell-surface F(1)-ATP synthase. FEBS J 277, 2192–2205 (2010).

13 Manalo, K. B., Choong, P. F., Becerra, S. P. & Dass, C. R. Pigment epithelium-derived factor as an anticancer drug and new treatment methods following the discovery of its receptors: a patent perspective. Expert Opin Ther Pat 21, 121–130 (2011).

14 Xi, L. Pigment Epithelium-Derived Factor as a Possible Treatment Agent for Choroidal Neovascularization. Oxid Med Cell Longev 2020, 8941057 (2020).

15 Qin, X. et al. PEDF is an antifibrosis factor that inhibits the activation of fibroblasts in a bleomycin-induced pulmonary fibrosis rat model. Respir Res 23, 100 (2022).

16 Nicolini, C. & Fahnestock, M. The valproic acid-induced rodent model of autism. Exp Neurol 299, 217–227 (2018).

17 Takuma, K. et al. Chronic treatment with valproic acid or sodium butyrate attenuates novel object recognition deficits and hippocampal dendritic spine loss in a mouse model of autism. Pharmacol Biochem Behav 126, 43–49 (2014).

18 Hara, Y. et al. Prenatal exposure to valproic acid increases miR-132 levels in the mouse embryonic brain. Mol Autism 8, 33 (2017).

19 Kawada, K., Mimori, S., Okuma, Y. & Nomura, Y. Involvement of endoplasmic reticulum stress and neurite outgrowth in the model mice of autism spectrum disorder. Neurochem Int 119, 115–119 (2018).

20 Traetta, M. E. et al. Hippocampal neurons isolated from rats subjected to the valproic acid model mimic in vivo synaptic pattern: evidence of neuronal priming during early development in autism spectrum disorders. Mol Autism 12, 23 (2021).

21 Moldrich, R. X. et al. Inhibition of histone deacetylase in utero causes sociability deficits in postnatal mice. Behav Brain Res 257, 253–264 (2013).

22 Kataoka, S. et al. Autism-like behaviours with transient histone hyperacetylation in mice treated prenatally with valproic acid. The international journal of neuropsychopharmacology 16, 91–103 (2013).

23 Kawanai, T. et al. Prenatal Exposure to Histone Deacetylase Inhibitors Affects Gene Expression of Autism-Related Molecules and Delays Neuronal Maturation. Neurochem Res 41, 2574–2584 (2016).

24 Liu, H. et al. Valproic Acid Induces Autism-Like Synaptic and Behavioral Deficits by Disrupting Histone Acetylation of Prefrontal Cortex ALDH1A1 in Rats. Front Neurosci 15, 641284 (2021).

25 Kundakovic, M., Chen, Y., Guidotti, A. & Grayson, D. R. The reelin and GAD67 promoters are activated by epigenetic drugs that facilitate the disruption of local repressor complexes. Mol Pharmacol 75, 342–354 (2009).

26 Tseng, C. J., McDougle, C. J., Hooker, J. M. & Zurcher, N. R. Epigenetics of Autism Spectrum Disorder: Histone Deacetylases. Biol Psychiatry 91, 922–933 (2022).

27 Maiti, A. et al. Class I histone deacetylase inhibitor suppresses vasculogenic mimicry by enhancing the expression of tumor suppressor and anti-angiogenesis genes in aggressive human TNBC cells. Int J Oncol 55, 116–130 (2019).

28 Blazejewski, S. M. et al. Rpsa Signaling Regulates Cortical Neuronal Morphogenesis via Its Ligand, PEDF, and Plasma Membrane Interaction Partner, Itga6. Cereb Cortex (2021).

29 Martinez-Cerdeno, V. Dendrite and spine modifications in autism and related neurodevelopmental disorders in patients and animal models. Dev Neurobiol 77, 393–404 (2017).

30 Mahmood, U. et al. Dendritic spine anomalies and PTEN alterations in a mouse model of VPA-induced autism spectrum disorder. Pharmacol Res 128, 110–121 (2018).

31 Baron-Mendoza, I. et al. Changes in the Number and Morphology of Dendritic Spines in the Hippocampus and Prefrontal Cortex of the C58/J Mouse Model of Autism. Front Cell Neurosci 15, 726501 (2021).

32 Risher, W. C., Ustunkaya, T., Singh Alvarado, J. & Eroglu, C. Rapid Golgi analysis method for efficient and unbiased classification of dendritic spines. PloS one 9, e107591 (2014).

33 Schneider, T. & Przewlocki, R. Behavioral alterations in rats prenatally exposed to valproic acid: animal model of autism. Neuropsychopharmacology 30, 80–89 (2005).

34 Dendrinos, G., Hemelt, M. & Keller, A. Prenatal VPA Exposure and Changes in Sensory Processing by the Superior Colliculus. Front Integr Neurosci 5, 68 (2011).

35 Mony, T. J., Lee, J. W., Dreyfus, C., DiCicco-Bloom, E. & Lee, H. J. Valproic Acid Exposure during Early Postnatal Gliogenesis Leads to Autistic-like Behaviors in Rats. Clin Psychopharmacol Neurosci 14, 338–344 (2016).

36 Zhao, S. et al. Genetic Testing Experiences Among Parents of Children with Autism Spectrum Disorder in the United States. J Autism Dev Disord 49, 4821–4833 (2019).

37 Martin, H. G. & Manzoni, O. J. Late onset deficits in synaptic plasticity in the valproic acid rat model of autism. Front Cell Neurosci 8, 23 (2014).

38 Wang, R. et al. Aberrant Development and Synaptic Transmission of Cerebellar Cortex in a VPA Induced Mouse Autism Model. Front Cell Neurosci 12, 500 (2018).

39 Soria-Ortiz, M. B., Reyes-Ortega, P., Martinez-Torres, A. & Reyes-Haro, D. A Functional Signature in the Developing Cerebellum: Evidence From a Preclinical Model of Autism. Front Cell Dev Biol 9, 727079 (2021).

40 Loo, L. et al. Single-cell transcriptomic analysis of mouse neocortical development. Nat Commun 10, 134 (2019).

41 Fombonne, E. Epidemiology of pervasive developmental disorders. Pediatr Res 65, 591–598 (2009).

42 Jeon, S. J. et al. Sex-specific Behavioral Features of Rodent Models of Autism Spectrum Disorder. Exp Neurobiol 27, 321–343 (2018).

43 Melancia, F. et al. Sex-specific autistic endophenotypes induced by prenatal exposure to valproic acid involve anandamide signalling. Br J Pharmacol 175, 3699–3712 (2018).

44 Kazlauskas, N., Seiffe, A., Campolongo, M., Zappala, C. & Depino, A. M. Sex-specific effects of prenatal valproic acid exposure on sociability and neuroinflammation: Relevance for susceptibility and resilience in autism. Psychoneuroendocrinology 110, 104441 (2019).

45 Ghahremani, R., Mohammadkhani, R., Salehi, I., Karimi, S. A. & Zarei, M. Sex Differences in Spatial Learning and Memory in Valproic Acid Rat Model of Autism: Possible Beneficial Role of Exercise Interventions. Front Behav Neurosci 16, 869792 (2022).

46 Weinstein-Fudim, L. et al. Gender Related Changes in Gene Expression Induced by Valproic Acid in A Mouse Model of Autism and the Correction by S-adenosyl Methionine. Does It Explain the Gender Differences in Autistic Like Behavior? Int J Mol Sci 20 (2019).

47 Notari, L. et al. Identification of a lipase-linked cell membrane receptor for pigment epithelium-derived factor. J Biol Chem 281, 38022–38037 (2006).

48 Bernard, A. et al. Laminin receptor involvement in the anti-angiogenic activity of pigment epithelium-derived factor. J Biol Chem 284, 10480–10490 (2009).

49 Subramanian, P., Notario, P. M. & Becerra, S. P. Pigment epithelium-derived factor receptor (PEDF-R): a plasma membrane-linked phospholipase with PEDF binding affinity. Adv Exp Med Biol 664, 29–37 (2010).

50 Park, K. et al. Identification of a novel inhibitor of the canonical Wnt pathway. Mol Cell Biol 31, 3038–3051 (2011).

51 Sagheer, U., Gong, J. & Chung, C. Pigment Epithelium-Derived Factor (PEDF) is a Determinant of Stem Cell Fate: Lessons from an Ultra-Rare Disease. J Dev Biol 3, 112–128 (2015).

52 Chan, N., He, S., Spee, C. K., Ishikawa, K. & Hinton, D. R. Attenuation of choroidal neovascularization by histone deacetylase inhibitor. PloS one 10, e0120587 (2015).

53 Matsuda, T. & Cepko, C. L. Electroporation and RNA interference in the rodent retina in vivo and in vitro. Proc Natl Acad Sci U S A 101, 16–22 (2004).

54 Blazejewski, S. M., Bennison, S. A., Liu, X. & Toyo-Oka, K. High-throughput kinase inhibitor screening reveals roles for Aurora and Nuak kinases in neurite initiation and dendritic branching. Sci Rep 11, 8156 (2021).

55 Chen, T.-W. et al. Ultrasensitive fluorescent proteins for imaging neuronal activity. Nature 499, 295–300 (2013).

